# Acidocalcisome-like vacuoles constitute a feedback-controlled phosphate buffering system for the cytosol

**DOI:** 10.1101/2025.05.30.656997

**Authors:** Samuel Bru, Lydie Michaillat Mayer, Geun-Don Kim, Danye Qiu, Henning Jacob Jessen, Andreas Mayer

## Abstract

Cells experience strong variations in the consumption and availability of inorganic phosphate (P_i_). Since P_i_ is an essential macronutrient but excess P_i_ has negative impacts on nucleotide hydrolysis and metabolism, its concentration must be maintained in a suitable range. Conserved storage organelles, acidocalcisomes, provide this buffering function. We used acidocalcisome-like yeast vacuoles to study how such organelles are set up to for this task. Our combined in vitro and in vivo analyses revealed that their ATP-driven polyphosphate polymerase VTC converts cytosolic P_i_ into inorganic polyphosphates (polyP), which it transfers into the vacuole lumen. Luminal polyphosphatases immediately hydrolyse this polyP to establish a growing reservoir of vacuolar P_i_. Product inhibition by this P_i_ pool silences the polyphosphatases, caps P_i_ accumulation, and favours vacuolar polyP storage. Upon cytosolic P_i_ scarcity, the declining inositol pyrophosphate levels activate the vacuolar P_i_ exporter Pho91 to replenish cytosolic P_i_. In this way, acidocalcisome-like vacuoles constitute a feedback-regulated buffering system for cytosolic P_i_, which the cells can switch between P_i_ accumulation, P_i_ release, and high-capacity phosphate storage through polyP.

## Introduction

Cells actively manage the concentration of P_i_ in their cytosol because they must strike a balance between conflicting goals (Austin & Mayer, 2020). On the one hand, P_i_ is a product of nucleotide-hydrolysing reactions. Its concentration has a significant impact on the free energy that these reactions can provide to drive metabolism. Since, for his reason, excessive P_i_ concentrations might stall metabolism, we can expect that cytosolic P_i_ must remain limited. But P_i_ is also an essential macronutrient. As a major constituent of nucleic acids and phospholipids, and as an important modifier of proteins, carbohydrates and many metabolites, it is consumed in large quantities for anabolic reactions. This can lead to situations where progression of the S-phase of the cell cycle is limited by the P_i_ uptake capacity of the cells (Bru *et al*, 2016, 2017; Gillies *et al*, 1981; Neef & Kladde, 2003). Rapidly dividing cells, such as yeast cells, remedy this problem by maintaining phosphate stores in the form of inorganic polyP. PolyPs are chains of P_i_ linked through phosphoric anhydride bonds, which can be stored in acidocalcisome-like organelles (Urech *et al*, 1978; Okorokov *et al*, 1980). Acidocalcisomes are lysosome-related organelles that have been characterised by pioneering work mainly in Trypanosomes, but they have been isolated from a multitude of other organisms (Docampo *et al*, 2005). Their defining chemical features are their acidity, high concentrations of polyP, Ca, Mg, Zn, Fe, K, Na, and basic amino acids and they are implicated in osmoregulation and the homeostasis of phosphate and metals (Docampo *et al*, 2005; Girard-Dias *et al*, 2023; Docampo, 2024). Yeast vacuoles share all these properties and are hence good and easily accessible models for studying aspects of acidocalcisomal mechanisms, such as the synthesis and turnover of polyP (Dürr *et al*, 1979; Huber-Wälchli & Wiemken, 1979; Urech *et al*, 1978; Austin & Mayer, 2020). But they have an additional, lysosome-like face, enabling them to assume also hydrolytic degradative functions for the cells (Li & Kane, 2009). Therefore, we refer to them as acidocalcisome-like organelles.

The polyP stores of vacuoles are accessed by cells when P_i_ becomes limiting in the environment, or when they face a sudden enhanced P_i_ consumption, e.g. during metabolic transitions from respiratory growth to fermentation, which requires much more phosphorylated metabolites (Shirahama *et al*, 1996; Thomas & O’Shea, 2005; Gillies *et al*, 1981; Nicolay *et al*, 1982, 1983). In such cases, cellular polyphosphate content decreases. It has therefore been proposed that polyPs are re-converted into P_i_ to buffer shortages in cytosolic P_i_ (Nicolay *et al*, 1983, 1982). While this hypothesis is straightforward, we are lacking a coherent concept of how a P_i_ buffer based on an acidocalcisome-like organelle might work. Pioneering work on these organelles has revealed several characteristic and conserved features, such as their acidity, and their high content of basic amino acids and divalent cations (Docampo & Huang, 2016; Docampo, 2024). Of direct relevance to P_i_ homeostasis is the high capacity of acidocalcisome-like organelles for storing polyP. PolyP can greatly vary in length, from two to hundreds of phosphate units. The membrane of acidocalcisome-like organelles can carry polyP polymerases, such as VTC, and P_i_ transporters, such as Pho91 (Jimenez & Docampo, 2015; Hürlimann *et al*, 2007; Wang *et al*, 2015; Huang & Docampo, 2015; Gerasimaite & Mayer, 2016; Müller *et al*, 2003, 2002). VTC is a coupled polyP polymerase and translocase (Gerasimaite *et al*, 2014), which synthesizes polyP by transferring the γ-phosphate of cytosolic ATP onto an elongating polyP chain (Hothorn *et al*, 2009). PolyP is generated by a catalytic domain in the centre of the Vtc4 subunit of the complex (Hothorn *et al*, 2009). The activity of this subunit is controlled through the SPX domains of VTC (Wild *et al*, 2016), which may associate into a dimerized inactive state (Pipercevic *et al*, 2023). They can be released from this state through the action of inositol pyrophosphates, signalling molecules that accumulate when P_i_ is abundant in the cytosol and activate P_i_ storage in the form of polyP. The P_i_ state of the cytosol is communicated to VTC through a specific inositol pyrophosphate, 1,5-IP_8_ (Gerasimaite *et al*, 2017; Chabert *et al*, 2023).

The postulated direct channelling of polyP from the site of its synthesis through the vacuolar membrane (Gerasimaite *et al*, 2014) is facilitated by the structure of the VTC complex. The catalytic domain of VTC has its exit for polyP right at the entry of a polyP-conducting channel, which is formed through the transmembrane domains of the VTC complex itself (Liu *et al*, 2023; Guan *et al*, 2023; Müller *et al*, 2002, 2003; Gerasimaite *et al*, 2014). This channel was proposed to exist in an open and closed conformation and to be gated through polyP (Liu *et al*, 2023). PolyP translocation may be driven by the electrochemical potential across the membrane, which could move the highly negatively charged polyP chain through electrophoresis. This can explain why efficient polyP synthesis depends on proton pumps such as the V-ATPase or H^+^-pumping pyrophosphatases (Gerasimaite & Mayer, 2016; Gerasimaite *et al*, 2014; Freimoser *et al*, 2006; Lemercier *et al*, 2004).

The transporter Pho91 was proposed to export vacuolar P_i_ (Hürlimann *et al*, 2007). Several properties of yeast Pho91 and its homologs from other organisms are consistent with this view. Its ablation increases vacuolar P_i_ and polyP content, in yeast as well as in trypanosomes (Hürlimann *et al*, 2007; Jimenez & Docampo, 2015; Farofonova *et al*, 2023), and it weakly induces the phosphate starvation program in yeast (Pinson *et al*, 2004), suggesting that it may induce cytosolic P_i_ scarcity. Furthermore, patch-clamp analyses of Pho91-like channels from yeast, trypanosomes and plants have shown their P_i_ permeability and the dependence of their directionality on a pH gradient across the membrane (Potapenko *et al*, 2018, 2019; Wang *et al*, 2015). In the presence of a pH gradient, the Pho91 homolog from rice, OsSpx-MFS3, mediates P_i_ flux along the proton gradient, consistent with a function in exporting P_i_ from the acidic lumen of vacuoles (Wang *et al*, 2015) towards the cytosol.

Acidocalcisome-like organelles also contain polyphosphatases in their lumen (Lander *et al*, 2016; Gerasimaite & Mayer, 2016; Kulakovskaya *et al*, 2021; McCarthy & Downey, 2023), which can convert polyP back into P_i_. In baker’s yeast, two vacuolar polyphosphatases are known, Ppn1 and Ppn2 (Gerasimaite & Mayer, 2017; Sethuraman *et al*, 2001; Andreeva *et al*, 2019). This means that a chain of polyP, when being synthesized by VTC and arriving in the vacuolar lumen, is immediately exposed to hydrolytic enzymes that will degrade it. While this seems at first sight paradoxical, we explored the hypothesis that the co-existence of polyP-synthesizing and polyP-hydrolysing activities might be a key feature conveying to acidocalcisome-like organelles the capacity to buffer cytosolic P_i_. That these organelles have a critical role to play in this process is illustrated by observations in yeast, where artificial up- or down-regulation of vacuolar polyP synthesis suffices to drive the cytosol into a state of P_i_ starvation or P_i_ excess, respectively (Desfougères *et al*, 2016a). Furthermore, the presence of polyP reserves delays the activation of the transcriptional phosphate starvation response, the PHO pathway (Thomas & O’Shea, 2005). We hence explored the capacity of isolated yeast vacuoles to interconvert polyP and P_i_, and we characterized the roles played by the vacuolar polyphosphatases Ppn1 and Ppn2 and the vacuolar P_i_ transporter Pho91. Our observations can be combined with previous findings to yield a coherent model of how an acidocalcisome-like organelle can operate as a P_i_ buffer for the cytosol.

## Results

We explored the interplay of VTC, polyphosphatases and Pho91 in the accumulation of polyP and P_i_ inside vacuoles using an in vitro system with purified organelles. Vacuoles can be isolated in intact form when the cells are gently opened by enzymatic digestion of the cell wall and disruption of the cell membrane by low concentrations of DEAE-dextran (Dürr *et al*, 1975). Organelles isolated in this way can perform many of their normal cellular functions, such as membrane fusion, membrane fission and autophagy (Michaillat *et al*, 2012; Sattler & Mayer, 2000; D’Agostino & Mayer, 2019; Kunz *et al*, 2004). They also contain active polyP polymerase (VTC) and active polyphosphatases (Ppn1 and Ppn2) and they can synthesize and import polyP (Gerasimaite *et al*, 2014; Gerasimaite & Mayer, 2017; Gerasimaite *et al*, 2017).

### Vacuolar P_i_ accumulation depends on polyP synthesis

Purified vacuoles were incubated with an ATP-regenerating system under conditions that allow these organelles to synthesize polyP in vitro (Gerasimaite *et al*, 2014). After different periods of incubation, the organelles were sedimented and solubilized in detergent. Vacuolar accumulation of polyP was assayed through DAPI fluorescence and P_i_ was measured through malachite green. VTC is stimulated by a variety of inositol pyrophosphates, most efficiently by 1,5-InsP_8_ (Gerasimaite *et al*, 2014, 2017). Since 1,5-InsP_8_ is not commercially available, we used saturating concentrations of 5-InsP_7_ for our experiments (Pavlovic *et al*, 2015). This compound stimulates VTC with a higher EC_50_ but to the same maximal activity as 1,5-InsP_8_, and it was more readily available to us for routine experiments. Under the conditions used here, the organelles rapidly and efficiently produced polyP (Fig. 1), in line with previous results (Gerasimaite *et al*, 2014, 2017). This production was stimulated by 5-InsP_7_, reaching 0.7 nmol of phosphate units per µg of vacuolar protein and min. This means that already within 10 min, wildtype vacuoles produced a mass of polyP equivalent to their total protein content, indicating how efficiently these organelles synthesize polyP. This signal was entirely dependent on VTC, because it was not observed in a mutant lacking the catalytic subunit of VTC (*vtc4Δ*).

**Fig. 1:**
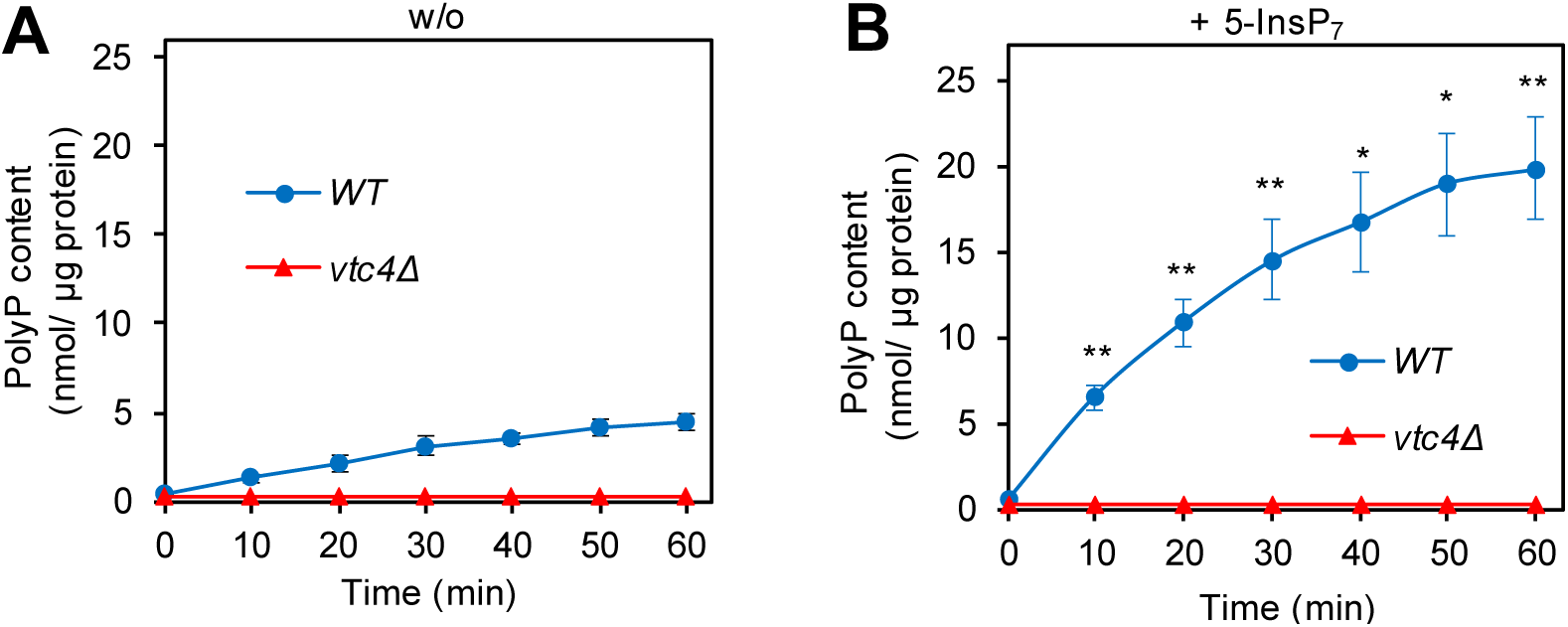
VTC- and 5-IP_7_-dependent polyP synthesis by isolated vacuoles. Vacuoles were isolated from logarithmic cultures of BY4742 wildtype cells (WT) or from isogenic vtc4Δ cells strains. They were incubated in polyphosphate synthesis assays without (*A*) or (B) in presence of 50 µM 5-InsP_7_. At indicated times, aliquots were withdrawn, solubilized in Triton X-100 and polyphosphate was quantified through the polyP-dependent fluorescence of added DAPI. Means ± SD (standard deviation) of three independent experiments are shown. * p<0.05, ** p<0.005 for a paired t-test comparing WT with and without 5-InsP_7_ at each time point.

Isolated wildtype vacuoles also accumulated P_i_ with significant efficiency, at an initial rate of at least 0.3 nmol µg^-1^ min^-1^ (Fig. 2A). Vacuoles from *vtc4Δ* cells showed only 8% of this P_i_ signal. Most of it was not time-dependent and hence may represent a background signal from the organelle preparation. In sum, vacuolar P_i_ accumulation depended on polyP synthesis through VTC.

**Figure 2.**
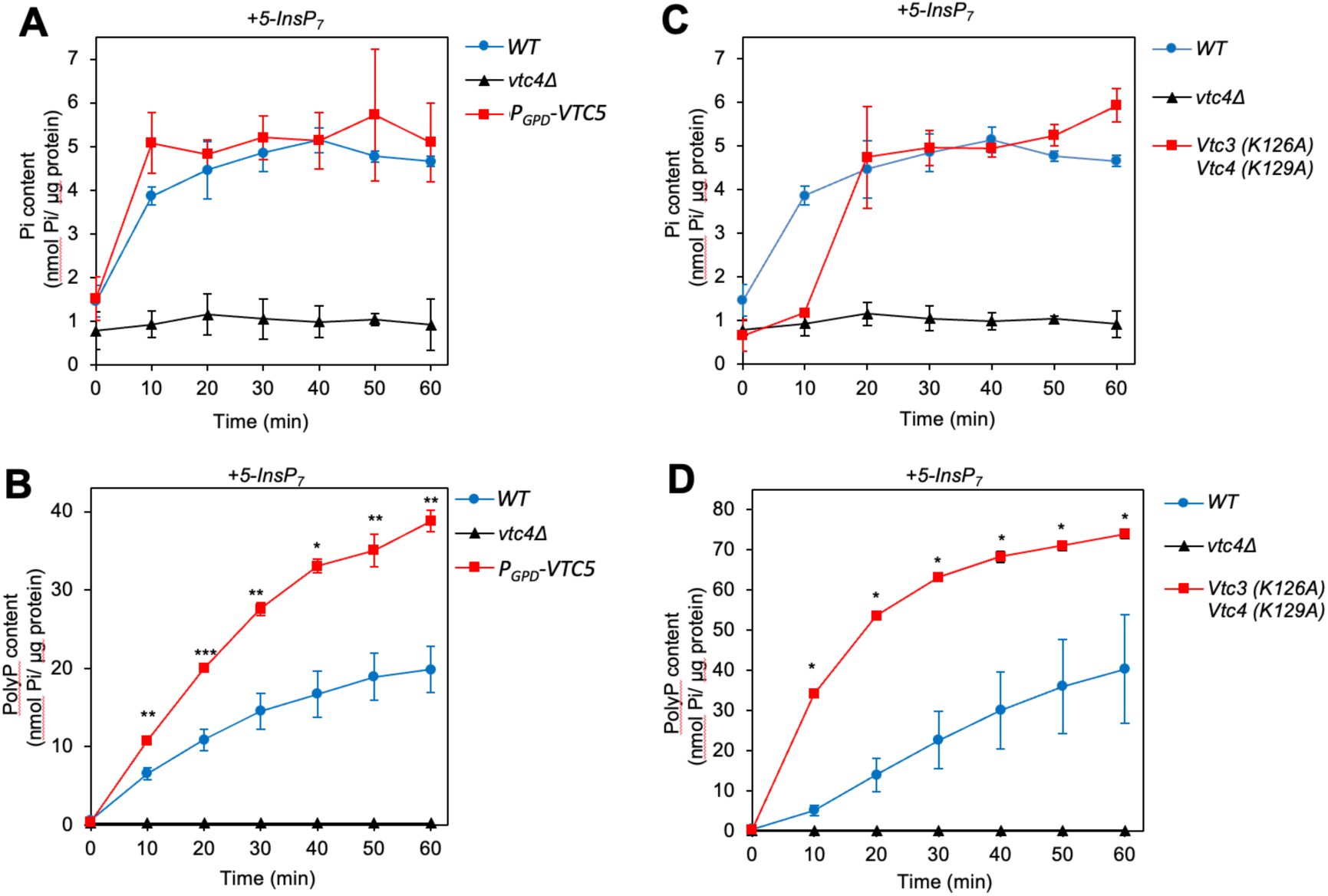
VTC-dependent accumulation of P_i_ in isolated vacuoles. Vacuoles were isolated from the indicated, logarithmically growing BY4742 (WT) cells. PolyP overproduction was achieved either through **(A, B)** overexpressing VTC5 from the strong GPD promotor, or **(C,D)** by expressing the hyperactivating vtc3^k126A^ and vtc4^K129A^ alleles form their native promotors as the sole source of these two proteins. The vacuoles were incubated with an ATP-regenerating system and 50 µM 5-InsP_7_ under conditions allowing polyP synthesis. At indicated times, the vacuoles were solubilized with Triton X-100 and P_i_ was assayed through malachite green. Graphs represent the mean ± SEM of three independent experiments for each strain. * p<0.05, ** p<0.005, *** p<0.001 from a paired t-test comparing each time WT vs VTC5 overexpression or vtc3^k126A^ vtc4^K129A^.

The kinetics of polyP and P_i_ accumulation were remarkably different. Whereas polyP continued to accumulate throughout the entire incubation period, P_i_ accumulated rapidly in the initial phase but reached a plateau within 30 min (Fig. 2A, B). Thus, although polyP production was essential for P_i_ accumulation, it could not become the limiting factor for it in the second phase. This is further illustrated by P_i_ accumulation in strains showing enhanced polyP synthesis, such as overexpressors of VTC5 (Fig. 2A, B) or strains carrying hyperactivating substitutions in the SPX domains of VTC (*vtc3^K126A^ vtc4^K129A^*; Fig. 2 C, D) (Desfougères *et al*, 2016b; Wild *et al*, 2016).

Vacuoles from these strains accumulated polyP at a two to seven times higher initial rate than the wildtype and to higher concentrations. Nevertheless, their accumulation of P_i_ arrested at the same level as that of wildtype vacuoles, at 5 nmol µg^-1^ vacuolar protein. To estimate the corresponding luminal concentration of P_i_, we measured the diameters of 100 isolated wildtype vacuoles and counted the number of vacuoles per µg of vacuolar protein. With an average diameter of 0.8 µm (Suppl. Fig. 1) and 5*10^7^ vacuoles/µg of protein we can estimate that P_i_ accumulation in vacuoles incubated with 5-InsP_7_ reached a plateau at a luminal concentration of around 30 mM. *vtc4Δ* vacuoles showed only a very small increase of less than 3 mM in luminal P_i_ content.

**Supplementary Figure 1:**
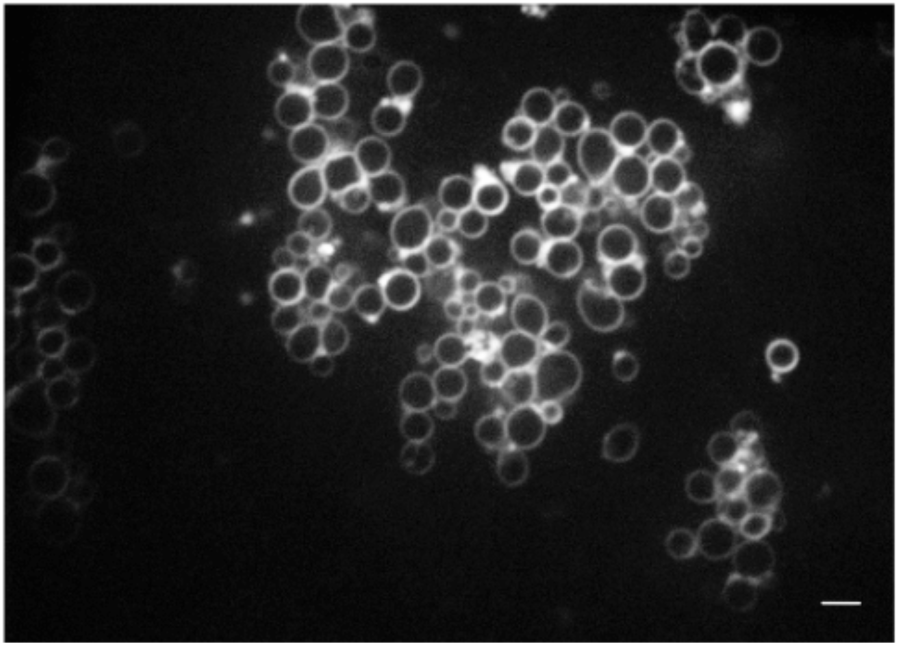
Isolated vacuoles during polyP synthesis. BJ 4742 (WT) vacuoles were incubated in vitro as in Fig. 1B. After 10 min, 0.1μM of the fluorophore FM4-64 was added to stain the membranes. The suspension was immediately imaged on a Yokogawa spinning disc microscope with a Gataca LiveSR unit and a 100x 1.45 NA objective. Stacks were taken at 0.2 µM z-distance. The image shows an optical section out of such a stack. To determine vacuole diameters, the stacks were subjected to maximum intensity projection in ImageJ. The largest visible diameter for 100 organelles was measured manually using ImageJ, yielding an average diameter of 0.8 µm with an SD of 0.2 µm Scale bar: 1 µm

The 30 mM of P_i_ accumulating in wildtype vacuoles is a substantial concentration, which is in the range of the in vivo P_i_ concentration of 25 mM that was measured by ^31^P-NMR spectroscopy of yeast cells under conditions where vacuolar P_i_ dominates the signal (Okorokov *et al*, 1980). Since, upon P_i_ scarcity, cytosolic P_i_ drops to 1 mM (Okorokov *et al*, 1980), a substantial P_i_ concentration gradient across the vacuolar membrane could drive rapid replenishment of the cytosol from a readily accessible vacuolar pool of P_i_.

### Vacuolar P_i_ accumulation is limited by feedback inhibition of Ppn1 and Ppn2

We asked why vacuolar P_i_ accumulation quickly forms a plateau whereas vacuolar polyP continues to accumulate. To this end, we explored the hypothesis that vacuolar polyphosphatases become inhibited when their product, P_i_, has accumulated. To test polyphosphatase activity we liberated the polyphosphatases from isolated vacuoles by detergent lysis, followed by incubation with synthetic polyP as a substrate and DAPI as an indicator for polyP. PolyP was efficiently degraded by this extract. The apparent polyPase activity was significantly reduced by 3 mM P_i_ and efficiently silenced by 30 mM P_i_ (Fig. 3A). To differentiate the contributions of Ppn1 and Ppn2, we also analysed the respective deletion mutants and performed the assay at higher polyP concentration in the presence Mg^2+^, which favours Ppn1 activity, or in the presence of Zn^2+^, which supports activity of Ppn1 and Ppn2 (Gerasimaite & Mayer, 2017). In the presence of Zn^2+^, the substrate was consumed in less than 3 min (Fig. 3). Degradation was delayed in vacuoles from *ppn1Δ* or *ppn2Δ* mutants, and it was suppressed in vacuoles from a *ppn1Δ ppn2Δ* mutant, in which both polyphosphatases were ablated. In incubations with only Mg^2+^ instead of Zn^2+^, which stimulates the activity of Ppn1 much more than that of Ppn2, polyP degradation was slower and genetic ablation of Ppn1 sufficed to stabilize polyP. The addition of 30 mM potassium phosphate, which is equivalent to the maximal P_i_ accumulation in vacuoles that we observed in Fig. 2, attenuated degradation of polyP 5-to 10-fold, both through Ppn1 (assayed in *ppn2Δ* and in samples without Zn^2+^) and through Ppn2 (assayed in *ppn1Δ*). That both polyphosphatases are inhibited by P_i_ at this concentration is consistent with the notion that product inhibition of Ppn1 and Ppn2 might limit the conversion of polyP into P_i_ in the vacuolar lumen and thus define the maximal concentration of the vacuolar P_i_ reservoir.

**Figure 3.**
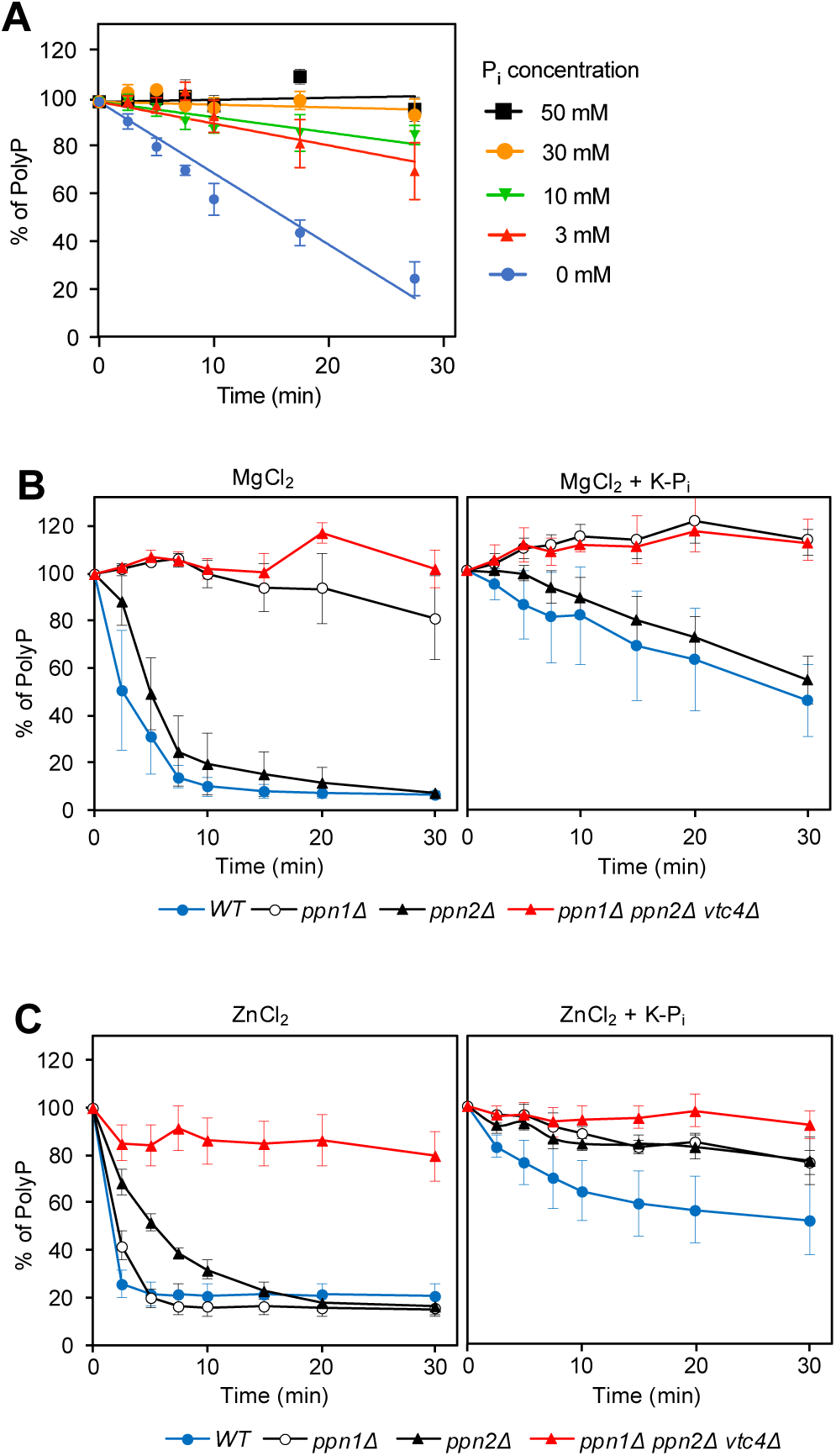
Effect of P_i_ on polyphosphatase activity. A. P_i_ titration. Vacuoles were isolated from logarithmically growing wildtype cells (BY4742). The organelles were diluted in polyphosphatase assay reaction buffer, which contained 0.1% Triton X-100 and hence liberated the luminal polyphosphatases. This lysate was incubated with 30 μM polyP_300_ as a substrate and supplemented with 1 mM ZnCl_2_ and the indicated concentrations of K-P_i_ pH 6.8. After the indicated times of incubation, the remaining polyP was quantified through DAPI. The DAPI signal at the beginning of the incubation served as 100% reference. Graphs represent the means ± SEM of three independent experiments. **B, C.** Differentiation of P_i_ effects on Ppn1 and Ppn2. Vacuoles were isolated from the indicated, logarithmically growing strains, lysed and used in polyP degradation assays with 300 µM polyP_300_ as in (A). These assays were performed in the presence or absence of 30 mM K-P_i_ pH 6.8 and 1 mM MgCl_2_ (B) or, instead of this, with 1 mM ZnCl_2_ (C) as cation supporting catalytic activity.

### Pho91 limits vacuolar P_i_ accumulation in an inositol pyrophosphate-dependent manner

To allow vacuoles to function as a P_i_ buffer for the cytosol, the P_i_ in the vacuolar lumen should become accessible in a regulated manner. The P_i_ transporter Pho91 is a prime candidate for mediating regulatable efflux because it is regulated by InsPPs through its SPX domain (Potapenko *et al*, 2018, 2019; Wang *et al*, 2015; Hürlimann *et al*, 2007). Furthermore, an overexpressed GFP fusion was reported to localize to vacuoles (Hürlimann *et al*, 2007; Elbaz-Alon *et al*, 2014), a phenotype that we confirmed (not shown). Since overexpression of membrane proteins in yeast easily leads to their erroneous accumulation in vacuoles, we re-investigated Pho91 localization in the absence of a strong, overexpressing promotor. When we expressed C- or N-terminal Pho91-GFP fusions from the authentic PHO91 promotor, we observed different and complex localization patterns (see Supplementary Figure 2 for examples).

**Supplementary Figure 2:**
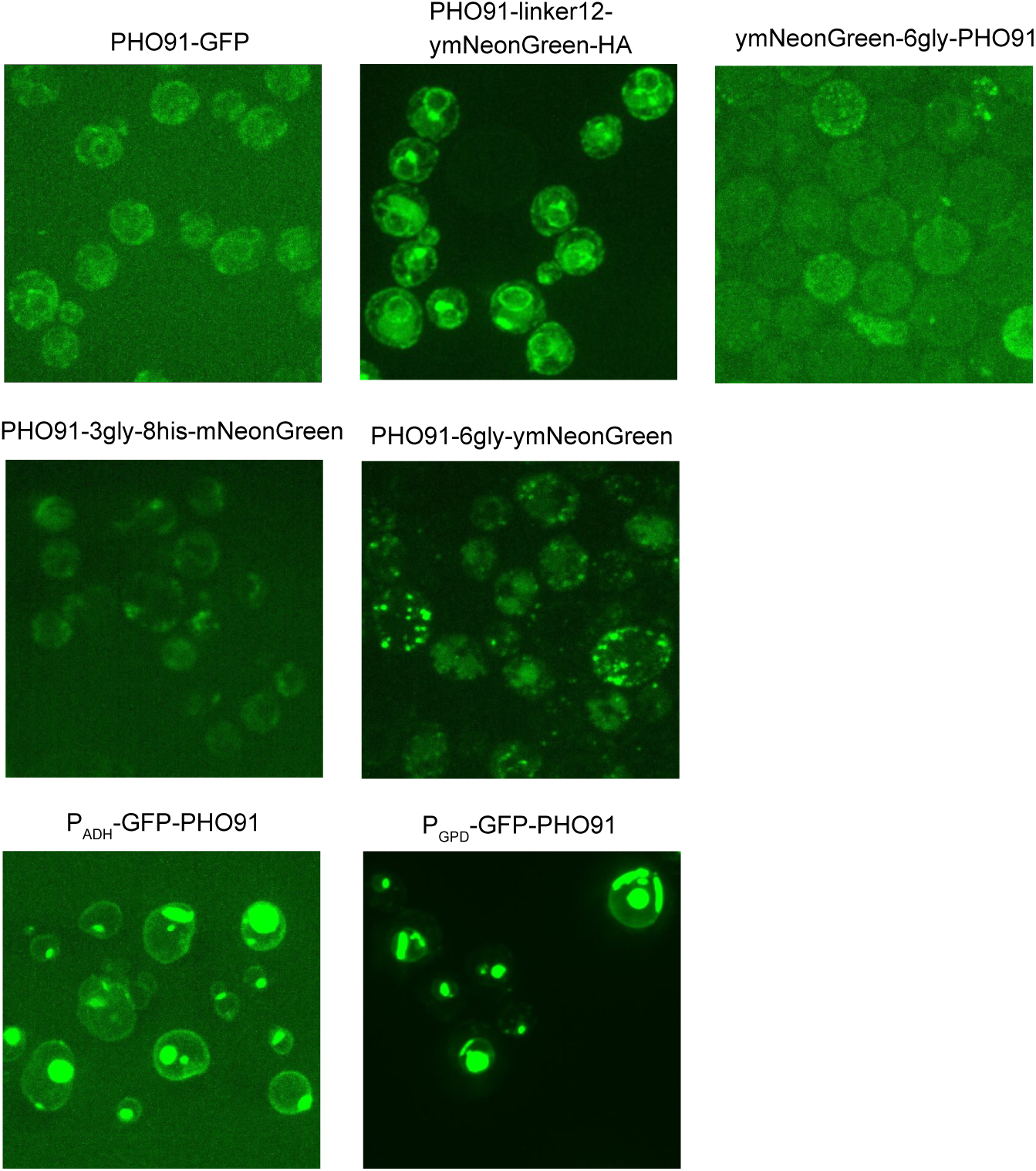
Effects of N- and C-terminal fluorescent protein tags on the localization of Pho91. PHO91 was fused with a variety of N- or C-terminal protein tags and peptide spacers as indicated. They were expressed from the endogenous PHO91 promotor or, where indicated, from the overexpressing ADH1 or GPD1 promotors. Cells were logarithmically grown overnight in SC medium, harvested at OD_600nm_ of 1-2, and analysed by fluorescence microscopy.

These localization patterns depended on the nature of the fluorescent tag and the linker peptides used to attach it to Pho91, and they often showed significant accumulation in the ER. This suggests that the fluorescent protein tags interfere with intracellular trafficking of Pho91. We hence analysed purified vacuoles to test whether the non-tagged, endogenous Pho91 is a vacuolar protein. Mass spectrometry was used to determine the enrichment of proteins in the vacuolar fraction relative to a whole cell extract (Suppl. Table 1). Peptides from Pho91 were enriched in the purified vacuoles to a similar degree (39-fold) as peptides from vacuolar marker proteins, such as the vacuolar polyP polymerase subunit Vtc3 (35-fold), the vacuolar amino acid transporter Avt3 (48-fold), the v-ATPase subunit Vma9 (46-fold), or the alkaline phosphatase Pho8 (34-fold). By contrast, typical plasma membrane proteins were barely enriched, such as the iron permease Ftr1 (2.5-fold), the P_i_ importer Pho87 (2.4-fold) or the polyamine importer Tpo5 (2.2-fold) (Supplementary data file 1). This co-enrichment suggests that the major fraction of non-tagged Pho91 indeed resides in the vacuole. In agreement with this, Pho91 engages the AP3-dependent vesicular trafficking pathway, which leads from the Golgi to the vacuole (Eising *et al*, 2022). In contrast to the other four yeast Pi transporters, Pho84, Pho87, Pho89, and Pho90, which individually suffice to support vigorous growth of yeast, Pho91 was reported to do support slower but significant growth and to fully rescue upon overexpression (Wykoff & O’Shea, 2001; Wykoff *et al*, 2007). Since this capacity to feed the cells is difficult to reconcile with a vacuolar localisation, unless the cells would feed through fluid phase endocytosis and subsequent export from the vacuole (Klompmaker *et al*, 2017), we reinvestigated this aspect. Instead of relying a down-regulatable PHO84 background, which results in a low but constitutive background expression of the potent Pi importer Pho84 in addition to Pho91, we relied on a more stringent approach, using plasmid shuffling to completely remove PHO89 when PHO91 is brought in. To this end, we generated a quintuple knockout strain lacking these five P_i_ transporters (Pho84, Pho87, Pho89, Pho90, Pho91), which was kept alive through expression of Pho89 from a URA-based plasmid. Plasmid shuffling allowed to exchange this plasmid against others expressing an individual P_i_ transporter. Whereas an individual plasma-membrane P_i_ transporter such as Pho89 supported normal colony formation, confirming previous observations (Wykoff *et al*, 2007; Wykoff & O’Shea, 2001), we could not confirm that Pho91 supports slower growth. In our plasmid shuffling approach, Pho91 did not support growth, at all (Suppl. Fig. 3). This supports that Pho91 is indeed a vacuolar P_i_ transporter, but that C- or N-terminal protein fusions interfere with correct sorting and cannot serve as reliable reporters for localization of this protein.

**Supplementary Figure 3:**
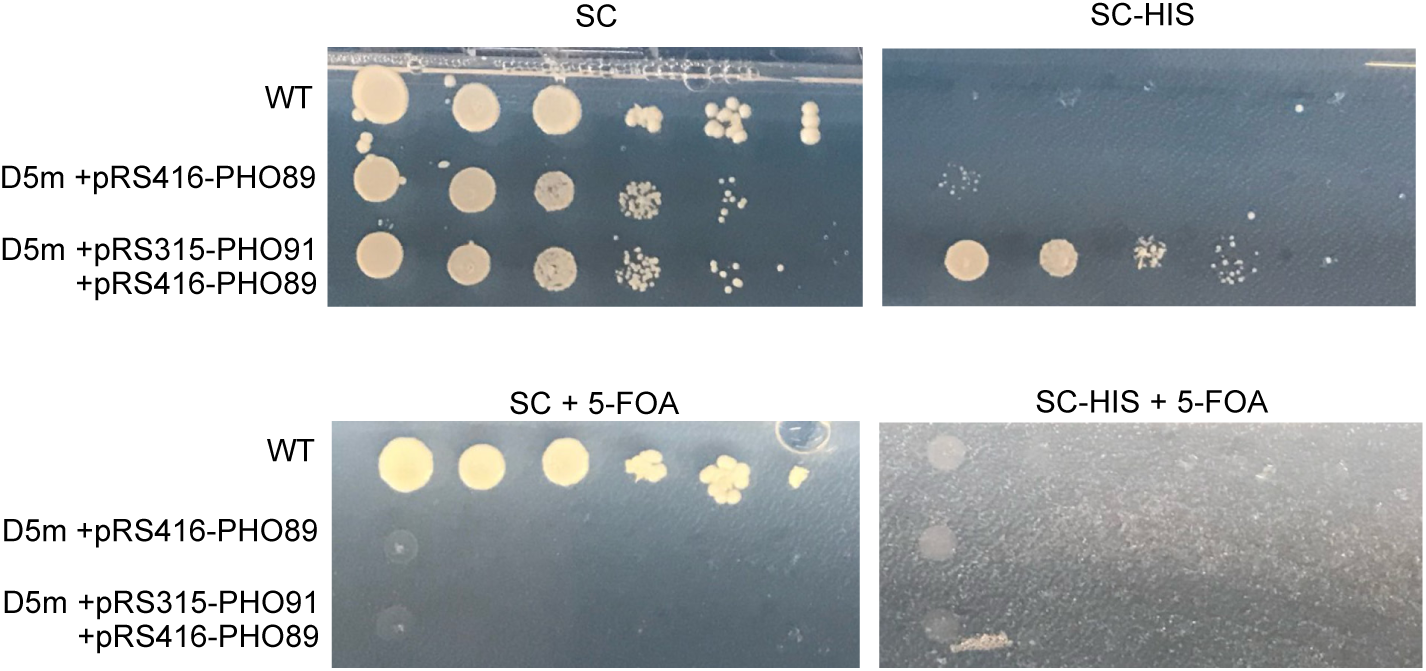
Pho91 cannot replace other P_i_ transporters to support growth of yeast. We generated a BY4741 strain with a quintuple deletion of the genes for the known P_i_ transporters, PHO84, PHO87, PHO89, PHO90 and PHO91 (D5m). These cells were kept alive by expressing the gene for the plasma membrane P_i_ transporter PHO89 from a URA3-based centromeric (single copy) plasmid (pRS416). PHO91 was expressed from a HIS3-based centromeric plasmid (pRS315). Cells were plated in a dilution series on SC lacking histidine (SC-HIS) to verify that the cells had the HIS3-based PHO91 plasmid, or on SC with 5-fluoro-orotic acid (5-FOA), a drug that forces cells to lose the URA3-based pRS416 and thus to live without PHO89. PHO91 as the sole P_i_ transporter gene (on SC + 5-FOA) does not allow cells to grow.

We tested the impact of Pho91 on P_i_ accumulation by vacuoles in vitro, using the same approach as above (Fig. 4). Vacuoles lacking Pho91 (*pho91Δ*) accumulated P_i_ two times faster than the wildtype and the maximal accumulated concentration was two times higher. This is consistent with a function of Pho91 as vacuolar P_i_ exporter (Hürlimann *et al*, 2007; Potapenko *et al*, 2018; Wang *et al*, 2015).

**Fig. 4:**
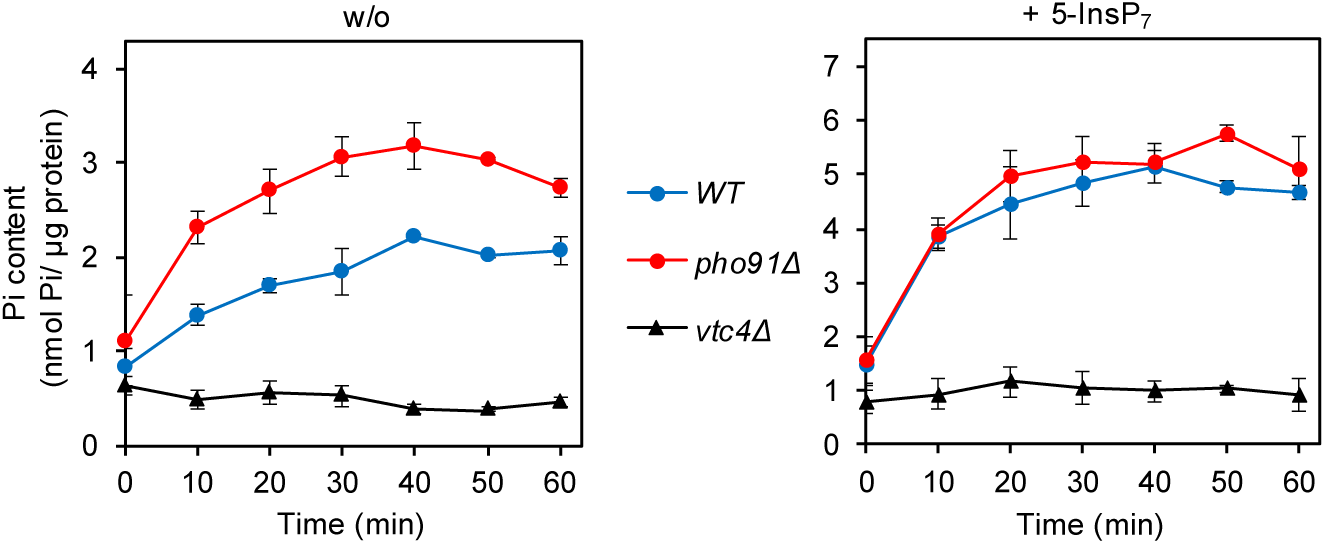
Accumulation of P_i_ in isolated vacuoles. VTC- and Pho91-dependence. Vacuoles were isolated from the indicated, logarithmically growing strains. The purified organelles were incubated as in Fig. 1, i.e. in a buffer with an ATP-regenerating system that allows the synthesis of polyP, and either (**A)** without (*w/o*) or (**B)** in the presence of 50 µM 5-IP_7_. After the indicated periods of incubation at 27°C, an 80 µl aliquot was withdrawn, the vacuoles were sedimented by centrifugation, washed and lysed. Released vacuolar P_i_ was determined by malachite green assay. Graphs represent the mean ± SEM of three independent experiments. p<0.05 from a paired t-test comparing each time WT versus *pho91Δ*.

The difference between *pho91Δ* and wildtype vacuoles vanished when the vacuoles were incubated in the presence of the inositol pyrophosphate 5-InsP_7_. Since, as we showed above, vacuolar P_i_ accumulation depends on polyP synthesis through VTC, the relative enhancement of P_i_ accumulation in wildtype vacuoles through 5-InsP_7_ could be caused by downregulation of Pho91, which would make the wildtype vacuoles behave similarly as *pho91Δ* vacuoles. Alternatively, 5-InsP_7_ might stimulate polyP synthesis in wildtype more than in *pho91Δ* vacuoles. We could rule out the latter explanation based on two observations: An assay of polyP synthesis during the incubation (Fig. 5) revealed that 5-InsP_7_ stimulated polyP synthesis in wildtype and *pho91Δ* vacuoles to similar degrees. Furthermore, as shown above (Fig. 2), polyP synthesis activity in wildtype cells is not rate-limiting for vacuolar P_i_ accumulation. Therefore, we attribute the enhancement of P_i_ accumulation through 5-InsP_7_ to Pho91. Upon P_i_ scarcity, the declining inositol pyrophosphate levels should then activate Pho91 to replenish cytosolic P_i_.

**Fig. 5:**
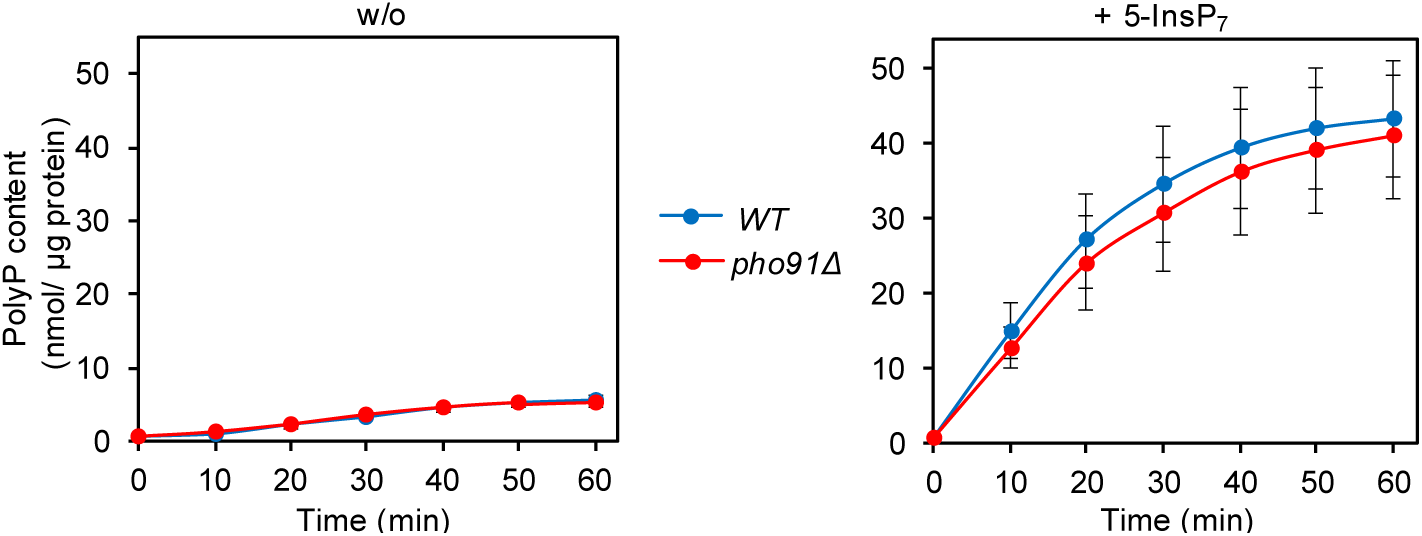
PolyP accumulation in pho91 mutant vacuoles. Vacuoles from BY4742 cells (WT) and isogenic *pho91Δ* cells were isolated and incubated under conditions supporting polyP synthesis and P_i_ accumulation as in Fig. 4, in the absence (w/o) or presence of 50 µM 5-InsP_7_. At the indicated timepoints, aliquots were withdrawn, the vacuoles were lysed in detergent, and polyP was assayed through DAPI fluorescence. Graphs show the means and SEM from three independent experiments. p>0.05 from a paired t-test for all differences between WT and *pho91Δ*.

### Interference with vacuolar polyP turnover provokes cytosolic P_i_ scarcity

We tested the effects of this postulated cycle of polyP synthesis, polyphosphatase activity and Pho91-mediated P_i_ export on P_i_ homeostasis in vivo. Pho4-GFP was used as a reporter, because this transcription factor shuttles between nucleus and cytosol. Under cellular P_i_ scarcity and correspondingly low inositol pyrophosphate levels it is predominantly nuclear, but it shifts to the cytosol under P_i_-replete conditions (O’Neill *et al*, 1996; Chabert *et al*, 2023; Desfougères *et al*, 2016a; Auesukaree *et al*, 2004). Pho4-GFP localisation can hence serve as a readout for cytosolic P_i_ signalling. In wildtype cultures growing logarithmically, cells can experience a transient shortage of P_i_ during S-phase, when P_i_ utilization for biosynthesis may exceed the uptake capacity of the cell and consume its polyP stores (Bru *et al*, 2016; Neef & Kladde, 2003; Pondugula *et al*, 2009). Therefore, even on P_i_-rich media, some cells can show nuclear accumulation of Pho4-GFP (Wykoff *et al*, 2007; Vardi *et al*, 2013, 2014). Since S-phase occupies only around 20% of the yeast cycle (Koren *et al*, 2010; Donaldson *et al*, 1998), this fraction is expected to be correspondingly small. However, it is significant, and it can be clearly quantified by microscopy, which scores individual cells and not the population as an ensemble. We exploited this natural fluctuation of intracellular P_i_ availability as a sensitising situation to assay how the vacuolar systems influence P_i_ balance in the cytosol.

We scored the fraction of logarithmically growing cells that show nuclear Pho4-GFP even in P_i_-replete medium. This fraction was 7% for wildtype cells (Fig. 6). It increased fourfold upon hyperactivation of polyP synthesis, which we achieved by overexpression of the regulatory VTC subunit Vtc5 (Desfougères *et al*, 2016a). Ablation of polyP synthesis by deletion of the catalytic subunit Vtc4 had the opposite effect and reduced the frequency of cells with nuclear Pho4-GFP by half. Deletion of the vacuolar polyphosphatases Ppn1 and Ppn2 prevents polyP turnover and leads to the accumulation of extremely long polyP chains (Gerasimaite & Mayer, 2017; Sethuraman *et al*, 2001). We may thus expect phosphate to remain fixed in the form of polyP instead of being made available for P_i_ reflux into the cytosol.

**Fig. 6:**
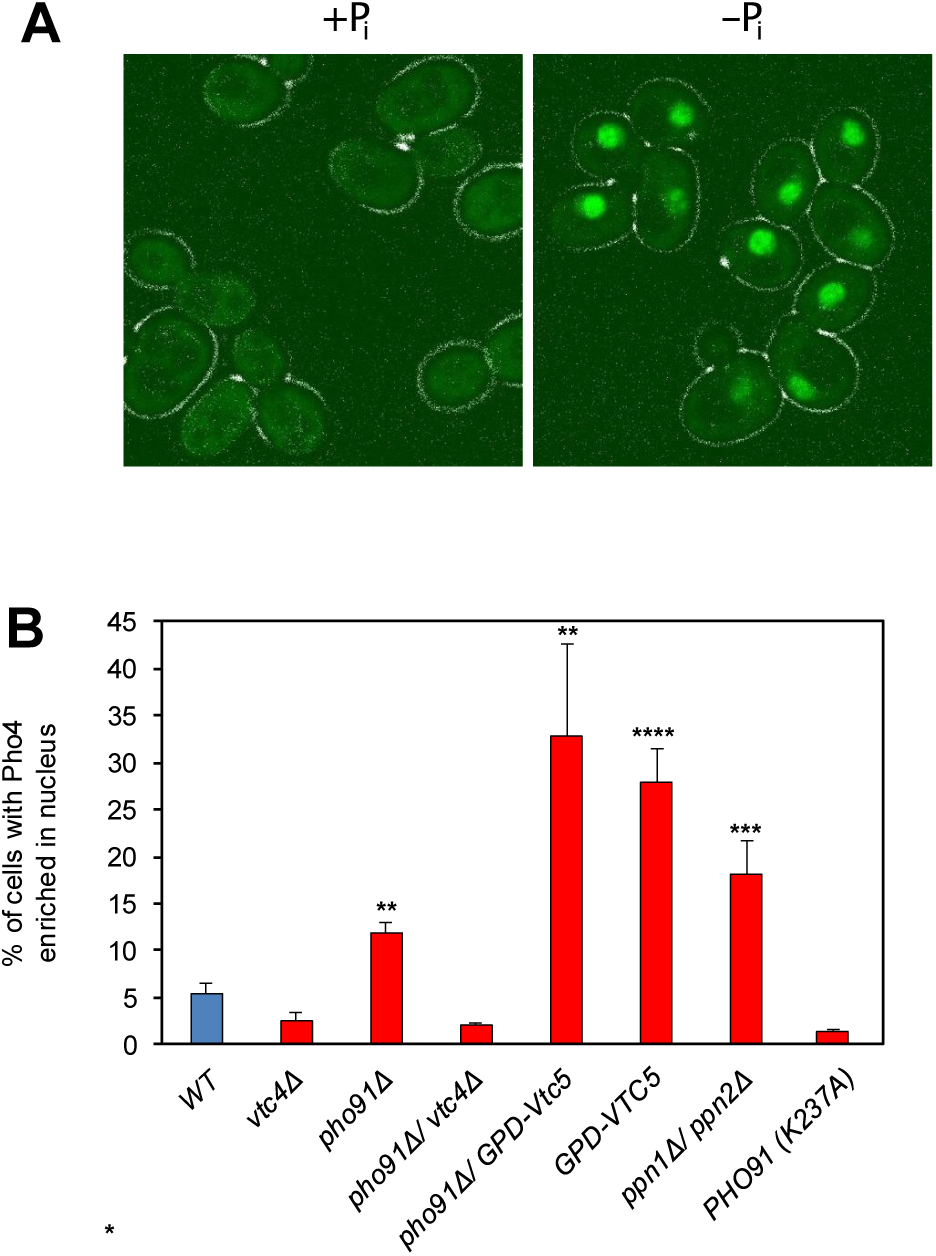
Impact of the vacuolar polyP/P_i_ cycle on cytosolic P_i_ signalling. A) Illustration of nucleo-cytoplasmic relocation of Pho4-GFP in response to P_i_ availability. Wildtype yeast cells were grown in SC medium under P_i_ replete conditions. During exponential phase (OD_600nm_=1), cells were transferred for 30 min to synthetic complete media with 200 µM phosphate (-P_i_) or 7.5 mM P_i_ (+P_i_) and imaged by fluorescence microscopy. B) The indicated yeast strains were logarithmically grown over night in SC medium with 7.5 mM P_i,_, harvested at OD_600nm_=1 and immediately imaged by fluorescence microscopy as in A. The graph shows the means and SEM of the percentage of cells showing Pho4 predominantly in the nucleus. n=3 independent experiments with 200 cells quantified per sample. ** p<0.005, *** p<0.001, ****p<0.0001 from a paired t-test comparing WT with the mutants.

In line with this, the fraction of *ppn1Δ ppn2Δ* double mutants that showed nuclear Pho4-GFP was 3-fold higher than in wildtype. *pho91Δ* cells, in which we expect P_i_ export from the vacuoles to be impaired, showed two times higher frequency of nuclear Pho4-GFP than wildtype. *pho91Δ vtc4Δ* double mutants showed an even lower frequency of nuclear Pho4-GFP than wildtype, suggesting that the state of P_i_ starvation that *pho91Δ* favours is dependent on vacuolar polyP accumulation. Cells expressing *pho91^K237A^* as the sole source of Pho91 also showed a 50% lower frequency of nuclear Pho4-GFP. The *pho91^K237A^* allele generates an amino acid substitution in the InsPP-binding patch of the Pho91 SPX domain. It mimics the InsPP-free state (Wild *et al*, 2016) and hence low-P_i_ conditions (Chabert *et al*, 2023). Collectively, our results are consistent with the notion that loss of InsPP binding activates Pho91 to export P_i_ from vacuoles to the cytosol, enhancing repression of the PHO pathway.

Since InsPP levels in yeast decline in response to cellular P_i_ availability (Lonetti *et al*, 2011; Chabert *et al*, 2023), we tested the impact of Pho91, Ppn1 and Ppn2 on the cellular levels of these metabolites using CE-MS, capillary electrophoresis coupled to mass spectrometry (CE-MS) (Qiu *et al*, 2020, 2023, 2021). The cells were harvested from the same P_i_-replete growth conditions as for the microscopic assays above (Fig. 7).

**Fig. 7:**
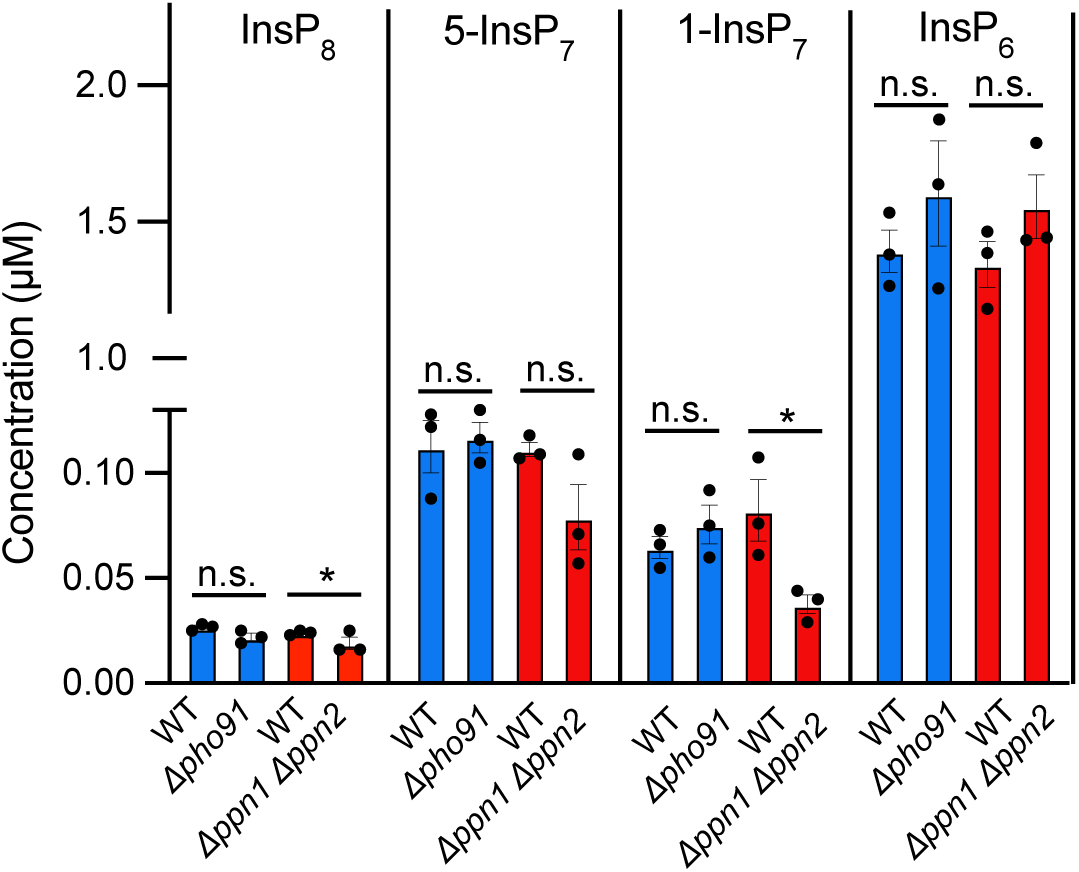
Impact of Pho91 and vacuolar polyphosphatases on InsPP levels. *pho91Δ* and *ppn1Δppn2Δ* cells, as well as their isogenic wildtypes, were logarithmically grown in P_i_-replete SC medium as in Fig. 6. At OD_600nm_=1 the cells were extracted with perchloric acid as previously described **(Wilson *et al*, 2015)** and analysed for the indicated InsPs through capillary electrophoresis coupled to mass spectrometry (CE-MS) as described **(Qiu *et al*, 2023)**. n=3. * p<0.05 from a paired t-test comparing WT with the mutants.

The *ppn1Δ ppn2Δ* double mutants showed a decrease by 30 to 50% in the inositol pyrophosphates 1,5-InsP_8_, 5-InsP_7_ and 1-InsP_7_, whereas InsP_6_ remained at a similar level as in wildtype. InsPP changes at this scale are functionally significant, because they suffice to trigger the initial phase of the P_i_ starvation response (Chabert *et al*, 2023; Kim *et al*, 2023, 2025). This is consistent with the 3-fold increase of cells with nuclear Pho4-GFP (Fig. 6). *pho91Δ* cells did not show significant changes for all four metabolites although they showed a partial shift of Pho4-GFP into the nucleus (Fig. 6). We attribute this discrepancy to the different nature of the assays. The microscopic assay for Pho4-GFP localization can pick up effects in a fraction of the cells because it offers single cell resolution. Inositol pyrophosphate analysis is an ensemble assay, in which changes in a smaller fraction of the population become diluted through the major pool that does not show the effect. For this reason, a change in inositol pyrophosphates affecting only 10% of the *pho91Δ* cells that show nuclear Pho4-GFP remains undetectable. The microscopic assay suggests that cytosolic P_i_ scarcity in *ppn1Δ ppn2Δ* affects more cells, perhaps because it is more profound, and it hence becomes detectable even in a whole-population-analysis.

## Discussion

Our observations can be integrated with existing data on the properties of VTC (Wild *et al*, 2016; Hothorn *et al*, 2009; Müller *et al*, 2002; Gerasimaite *et al*, 2014, 2017; Liu *et al*, 2023; Guan *et al*, 2023; Pipercevic *et al*, 2023) and Pho91 (Hürlimann *et al*, 2007; Wang *et al*, 2015; Potapenko *et al*, 2018) to generate a working model explaining how an acidocalcisome-like organelle such as the yeast vacuole is set up to function as a P_i_ buffer for the cytosol. Under P_i_-replete conditions, high InsPP levels activate VTC to polymerize P_i_ into polyP and translocate it into the vacuolar lumen. Here, the vacuolar polyphosphatases degrade polyP into P_i_, filling the lumen with P_i_ (Sethuraman *et al*, 2001; Shi & Kornberg, 2005; Lichko *et al*, 2010; Gerasimaite & Mayer, 2017). Since the P_i_ exporter Pho91 is downregulated through InsPP binding to its SPX domain (Potapenko *et al*, 2018; Hürlimann *et al*, 2007; Wang *et al*, 2015), the P_i_ liberated through polyP hydrolysis accumulates in the vacuoles. Product inhibition of the polyphosphatases attenuates polyP hydrolysis once the vacuolar lumen has reached a P_i_ concentration above 30 mM. When the cells experience P_i_ scarcity, InsPP levels decline (Chabert *et al*, 2023). This activates Pho91 to release P_i_ from the vacuolar pool into the cytosol and stabilizes cytosolic P_i_.

**Fig. 8:**
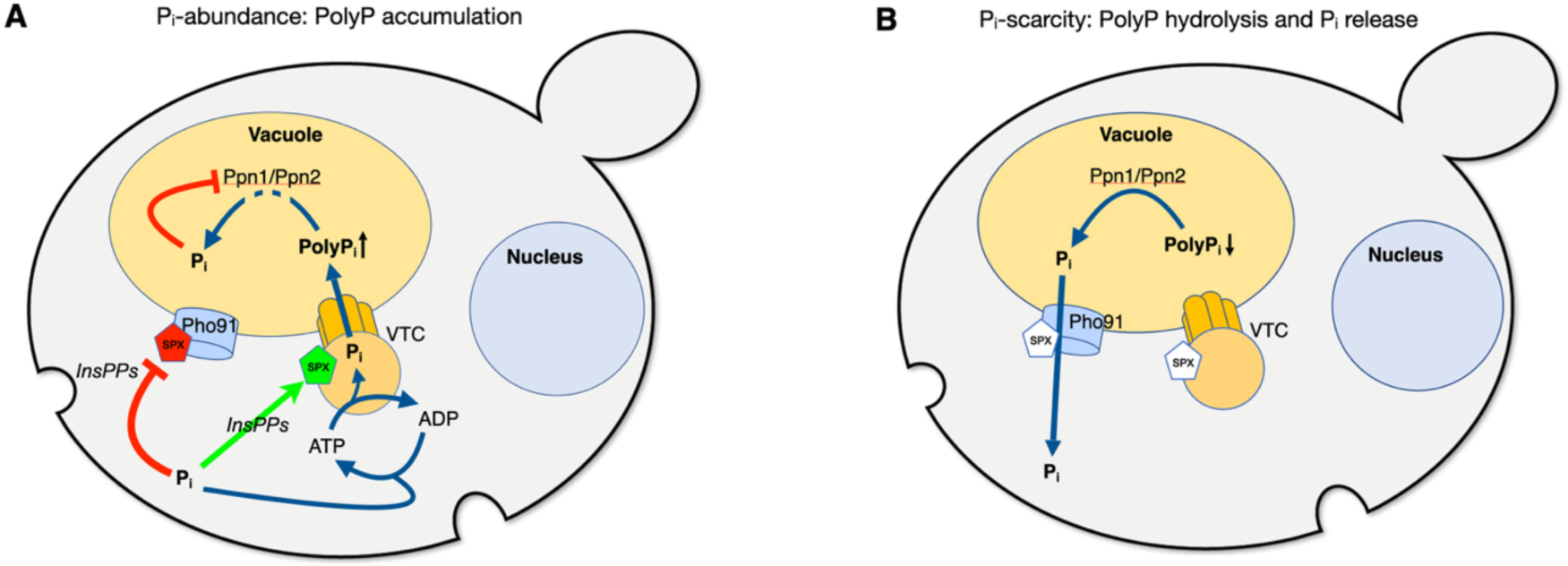
Working model of acidocalcisome-like vacuoles as P_i_ buffering systems. (A) Under Pi-replete conditions, ATP drives the conversion of P_i_ into polyP and its translocation into the organelle. Here, polyP is degraded by the vacuolar polyphosphatases Ppn1 and Ppn2 to establish a vacuolar pool of free P_i_. Feedback inhibition of P_i_ gradually reduces polyP degradation, enabling the buildup of a vacuolar polyP stock. Red lines and SPX colouring indicate inhibitory action, green colouring stimulation. (B) Cytosolic P_i_ scarcity decreases InsPP levels, which triggers two compensatory, SPX-controlled effects: The transfer of P_i_ from the cytosol into vacuoles through VTC ceases; and Pho91-dependent export of P_i_ from vacuoles is activated. Both measures synergize to stabilize cytosolic P_i_. The export of P_i_ from the vacuole lifts product inhibition on the polyphosphatases Ppn1 and Ppn2 and stimulates a compensatory degradation of polyP.

Yeast cells do not only accumulate P_i_ as a rapidly accessible buffer for the cytosol. Under P_i_-replete conditions they accumulate hundreds of millimolar of phosphate in the form of polyP (Urech *et al*, 1978). In contrast to the vacuolar reserve of P_i_, which is presumed to be accessible immediately, mobilizing the polyP store takes minutes to hours (Nicolay *et al*, 1982; Pondugula *et al*, 2009; Bru *et al*, 2016). But the polyP store offers advantages in the form of high capacity - hundreds of millimolar of phosphate units can be stored in the form of polyP - and low osmotic activity of polyP (Dürr *et al*, 1979). Keeping such a large stock of a critical resource, which is often growth-limiting in nature, is relevant for the cells. In case of phosphate shortage, the vacuolar polyP store can be mobilized to enable the cells to complete the cell cycle and transition into G_0_ phase (Spain *et al*, 2015; Westenberg *et al*, 1989; Müller *et al*, 1992). This can consume substantial amounts of phosphate, because we can estimate that replicating the entire DNA (1.2*10^7^ base pairs) immobilizes roughly 1 mM P_i_ in the cells; and cellular RNA is even 50 times more abundant than DNA (Warner, 1999), accounting for 50 mM phosphate. Phospholipids, which must also be synthesized to complete a cell cycle, fix phosphate in similar amounts (Lange & Heijnen, 2001). Thus, a large polyP store is necessary to guarantee that the cells can finish S-phase upon a shortage of phosphate sources. In accord with this notion, absence of the polyP store impairs cell cycle progression, nucleotide synthesis and induces genome instability (Bru *et al*, 2016, 2017). Also, a shift from non-fermentable carbon sources to fermentation of glucose leads to a strong requirement for P_i_ because the activation of glucose uptake and glycolysis depends on large amounts of phosphate-containing sugars and glycolytic intermediates (Nicolay *et al*, 1982, 1983; Gillies *et al*, 1981). Shortage of P_i_ restrains the abundance of these metabolites (Kim *et al*, 2023).

The properties of the regulatory circuit described above imply an inbuilt switch from vacuolar P_i_ accumulation to large-scale stocking of vacuolar polyP. P_i_-replete conditions generate high cellular InsPP levels. These will not only reduce P_i_ efflux from the vacuoles through Pho91 and inactivate the vacuolar polyphosphatases, but at the same time stimulate continued polyP synthesis by VTC. Coincidence of these effects will favour storage and high accumulation of phosphate in the form of polyP. Conversely, depletion of the vacuolar P_i_ reservoir upon P_i_ scarcity in the medium will activate the vacuolar polyphosphatases. In combination with the downregulation of the polyP polymerase VTC through the decline of InsPPs this will mobilize the large vacuolar polyP reserve once the immediately available vacuolar P_i_ pool is gradually depleted.

The concentration of P_i_ inside vacuoles as a rapidly accessible P_i_ reserve, and the synthesis of a large polyP stock, come at an energetic cost because the transformation of P_i_ into polyP requires the formation of phosphoric anhydride bonds (Hothorn *et al*, 2009; Gerasimaite *et al*, 2014) and vacuolar P_i_ reaches 30 mM. This exceeds the cytosolic P_i_ concentration, which was measured through ^31^P-NMR in a variety of yeasts, yielding values of 5-17 mM (Nicolay *et al*, 1982, 1983). Cytosolic P_i_ can also be estimated based on data from several other studies (Auesukaree *et al*, 2004; Hürlimann *et al*, 2009; Theobald *et al*, 1996; Zhang *et al*, 2015; Pinson *et al*, 2004). Assuming that the cytosolic volume of a BY4741 yeast cell is 40 fL (Chabert *et al*, 2023), and 1 g of dry weight contains 40 * 10^9^ yeast cells, these studies point to cytosolic values of 10-15 mM in P_i_-replete media. Upon P_i_ starvation, this value rapidly drops up to fivefold, resulting in a strong P_i_ gradient across the vacuolar membrane (Okorokov *et al*, 1980; Shirahama *et al*, 1996). To replenish the cytosolic pool under P_i_ scarcity, Pho91 can exploit not only this P_i_ concentration gradient, but also the vacuolar electrochemical potential, which was shown to stimulate P_i_ export through the Pho91 homolog OsSPX-MFS3 from plant vacuoles (Wang *et al*, 2015).

Vacuolar P_i_ accumulation is driven indirectly through ATP in two ways. VTC uses ATP as a substrate and transfers the phosphoric anhydride bond of the γ-phosphate onto a polyP chain (Hothorn *et al*, 2009). The growing polyP chain exits from the catalytic site directly towards the transmembrane part of VTC (Liu *et al*, 2023; Guan *et al*, 2023). This transmembrane part likely forms a controlled channel that can guide polyP through the membrane (Liu *et al*, 2023). Coupled synthesis and translocation require the V-ATPase (Gerasimaite *et al*, 2014), probably because polyP is highly negatively charged and therefore follows the electrochemical potential across the vacuolar membrane of 180 mV (inside positive) and 1.7 pH units (Kakinuma *et al*, 1981), which is generated through the proton pumping V-ATPase. Thus, the combination of the VTC complex and vacuolar polyphosphatases can be considered as a P_i_ pump that is driven by ATP through polyP synthesis and through the electrochemical potential for polyP translocation and P_i_ export.

It is likely that acidocalcisome- and lysosome-like organelles of other organisms act as buffers for cytosolic P_i_ similarly as described in our model for yeasts. This notion is supported by the conserved molecular setup of acidocalcisome-like organelles as well as by phenotypic similarities. The acidocalcisomes of Trypanosomes contain VTC, a Pho91 homolog and proton pumps in their membranes, and polyphosphatases in their lumen (Huang & Docampo, 2015; Billington *et al*, 2023; Fang *et al*, 2007; Lander *et al*, 2013; Ulrich *et al*, 2013; Scott *et al*, 1997). Also the acidocalcisome-like organelles of the alga Chlamydomonas contain such proteins and they accumulate polyP through VTC as a function of the availability of P_i_, a proton gradient and metal ions (Zúñiga-Burgos *et al*, 2024; Blaby-Haas & Merchant, 2014; Long *et al*, 2023; Hong-Hermesdorf *et al*, 2014; Aksoy *et al*, 2014; Ruiz *et al*, 2001; Goodenough *et al*, 2019). Like in yeast, the polyP stores are mobilized upon P_i_ limitation (Sanz-Luque *et al*, 2020; Plouviez *et al*, 2021). Drosophila has a potentially lysosome-related compartment, which is acidic, carries V-ATPase and a homolog of the Pi exporter XPR1, impacts cytosolic P_i_ and diminishes upon P_i_ starvation (Xu *et al*, 2023). Mammalian lysosome-like organelles also participate in P_i_ homeostasis. They can accumulate polyP and take up P_i_ (Pisoni, 1991; Pisoni & Lindley, 1992). They carry the P_i_ exporter XPR1, which interacts with the plasma membrane P_i_ importer PiT1 to regulate its degradation (Li *et al*, 2024).

We hence propose that acidocalcisome-like vacuoles may have a general role as feedback-controlled, rapidly accessible P_i_ buffers for the cytosol, addressing a critical parameter for metabolism. However, given that acidocalcisome-like organelles accumulate not only phosphate but also multiple other metabolites and ions (Docampo, 2024), they are probably interlinked with cellular metabolism in multiple ways and might form an important hub for its homeostasis.

## Materials and Methods

### Yeast strains and growth conditions

Saccharomyces cerevisiae cells were grown on yeast extract-peptone-dextrose (YPD: 1% yeast extract, 2% peptone and 2% dextrose) or in synthetic complete (SC) medium from Formedium, supplemented with sodium phosphate as needed). Yeast backgrounds used in this study were BY4741 and BY4742. Genetic manipulations of yeast were performed by homologous recombination according to published procedures and/or transformation with the indicated plasmids (Gietz & Schiestl, 2007; Güldener *et al*, 1996). Strains used are listed in Table 1.

**Table 1:**

| Background | Genotype | Source |
| --- | --- | --- |
| BY4741 | MATa his3Δ1 leu2Δ0 met15Δ0 ura3Δ0 | Euroscarf |
| BY4742 | MATα his3Δ1 leu2Δ0 lys2Δ0 ura3Δ0 | Euroscarf |
| BY4741 | *vtc4::kanMX* | (Müller *et al*, 2002) |
| BY4741 | *pho91::kanMX* | This study |
| BY4742 | *ppn1::natNT2* | (Gerasimaite & Mayer, 2017) |
| BY4742 | *ppn2::URA3* | Gerasimaite & Mayer, 2017) |
| BY4741 | VTC5 natNT2-GPD1pr | (Desfougères *et al*, 2016a) |
| BY4742 | VTC3<sup>K126A</sup> VTC4<sup>K129A</sup> | (Wild *et al*, 2016) |
| BY4741 | *pho91::kanMX* pRS303-P<sub>PHO91</sub>-PHO91 (K237A) | This study |
| BY4741 | *pho91::kanMX* pRS303-P<sub>PHO91</sub>-PHO91 (Y22F) | This study |
| BY4741 | *vtc4::kanMX pho91::hyg* | This study |
| BY4741 | pRS415-P<sub>PHO4</sub>-PHO4-GFP | This study |
| BY4741 | *vtc4::kanMX* pRS415-P<sub>PHO4</sub>-PHO4-GFP | This study |
| BY4741 | *pho91::kanMX* pRS415-P<sub>PHO4</sub>-PHO4-GFP | This study |
| BY4741 | *vtc4::kanMX pho91::hyg* pRS415-P<sub>PHO4</sub>-PHO4-GFP | This study |
| BY4741 | VTC5 natNT2-GPD1pr *pho91::hyg* pRS415-P<sub>PHO4</sub>-PHO4-GFP | This study |
| BY4741 | VTC5 natNT2-GPD1pr pRS415-P<sub>PHO4</sub>-PHO4-GFP | This study |
| BY4742 | *ppn1::natNT2 ppn2::URA3* pRS415-P<sub>PHO4</sub>-PHO4-GFP | This study |
| BY4741 | *pho91::kanMX* pRS303-P<sub>PHO91</sub>-PHO91 (K237A)<br>pRS415-P<sub>PHO4</sub>-PHO4-GFP | This study |
| BY4741 | pRS416-P<sub>ADH</sub>-SPL2 *pho91::kanMX* pRS303-P<sub>PHO91</sub>-PHO91 (K237A) pRS415-P<sub>PHO4</sub>-PHO4-GFP | This study |

### PHO4 localization

Cells transformed with pRS415-P_PHO4_-PHO4-GFP, a plasmid expressing a PHO4-GFP fusion from the PHO4 promotor (Chabert *et al*, 2023), were grown exponentially overnight in SC medium without leucine (SC-Leu). Care was taken that the culture did not grow beyond a density of OD_600_=0.7. At this point, the cells were collected by brief centrifugation (15 sec, 3000 x g) in a tabletop centrifuge and resuspended in 1/10th to 1/20th of their own supernatant. Pho4-GFP localization was immediately checked by microscopy. An aliquot of the cells was washed twice with SC-Leu without P_i_ and then diluted in SC-Leu with 200 μM phosphate to OD_600_=0.7. After 30 minutes of incubation in this medium, Pho4-GFP localization was analysed by fluorescence microscopy on a LEICA DMI6000B inverted microscope equipped with a Hamamatsu ORCA-R2 (C10600-10B) camera, an XCite ® series 120Q UV lamp and a Leica 100x 1.4 NA lens.

### Vacuole preparation

Vacuoles were purified from yeast cells essentially as described (D’Agostino & Mayer, 2019). Briefly, yeast cells were grown in 1L of YPD to an OD_600_ of 1.5. 330 ml of cells were collected by centrifugation and resuspended in 50 ml of 30 mM Tris-HCl pH 8.9, 10 mM DTT buffer. Suspensions were incubated in a 30° C water bath for 5 min and collected by centrifugation. The pellet was resuspended in 15 ml of spheroplasting solution (50 mM K-phosphate pH 7.5, 600 mM sorbitol in YPD with 0.2% D-glucose and 3600 U/ml lyticase) and incubated for 25 min at 30°C. Spheroplasts were collected by centrifugation (2500 x g, 3 min) and resuspended in 2.5 ml of 15% Ficoll 400 in PS buffer (10 mM PIPES-KOH pH 6.8, 200 mM sorbitol). 80 μg of DEAE-dextran were added under gentle mixing. After incubation on ice for 2 min and then at 30 °C for 80 sec, spheroplasts were transferred into Beckman SW41.1 tubes, overlaid with cushions of 8%, 4% and 0% Ficoll 400 in PS buffer, and centrifuged (150’000 x g, 90 min, 2 °C). Vacuoles were collected from the 0-4 % Ficoll interface. Their protein concentration was determined through Bradford assay using BSA as a standard.

### Polyphosphatase activity of vacuolar lysates

Polyphosphatase activity was assayed as described previously (Gerasimaite & Mayer, 2017), with the following modifications. Isolated vacuoles were diluted to a final protein concentration of 0.002 mg/ml in 1 ml of reaction buffer (20 mM PIPES/KOH pH 6.8, 150 mM KCl, 1 mM ZnCl_2_ or MgCwwl_2_, 0.1% Triton X-100, 1xPIC, 1 mM PMSF and 30 or 300 µM polyP_300_) in the presence or absence of various concentrations of KH_2_PO_4_ and incubated at 27°C. At the indicated times, 80 µl aliquots were collected and the reaction was stopped by dilution with 160 µl of stop solution (10 mM PIPES/KOH pH 6.8, 150 mM KCl, 12 mM EDTA pH 8.0, 0.1% Triton X-100, 15 µM DAPI). Remaining polyP was quantified by measuring polyP-DAPI fluorescence (λ_exc._ 415 nm, λ_em._ 550 nm) in a black 96 well plate in a Spectramax Gemini microplate fluorometer (Molecular Devices). A reaction containing boiled vacuoles was used as negative control.

### Polyphosphate synthesis by isolated vacuoles

Polyphosphate synthesis was assayed as described (Gerasimaite *et al*, 2014). Isolated vacuoles were diluted to final protein concentration of 0.02 mg/ml on 1 ml of reaction buffer (10 mM PIPES/KOH pH 6.8, 150 mM KCl, 0.5 mM MnCl2, 200 mM sorbitol) and the reaction was started by adding an ATP regenerating system (1 mM ATP-MgCl_2_, 40 mM creatine phosphate and 0.25 mg/ml creatine kinase). The mix was incubated at 27 °C. At different timepoints, 80 µl aliquots were mixed with 160 µl of stop solution (10 mM PIPES/KOH pH 6.8, 150 mM KCl, 200 mM sorbitol, 12 mM EDTA, 0.15% Triton X-100 and 15 μM DAPI). PolyP synthesis was quantified through polyP-DAPI fluorescence (λ_exc._ 415 nm, λ_em._ 550 nm) in a black 96 well plate. A calibration curve was prepared using commercial polyP_60_ as a standard.

### Phosphate quantification in isolated vacuoles

Isolated vacuoles were incubated as described for the polyP synthesis assay above. At different time points, 80 µl aliquots were centrifuged (3 min, 2000 x g, 2°C), the pellets were washed with 500 µl of washing solution (10 mM PIPES/KOH pH 6.8, 200 mM Sorbitol, 150 mM KCl) and centrifuged as before. The final pellet was resuspended with 100 µl of lysis buffer (10 mM PIPES/KOH pH 6.8, 200 mM sorbitol, 150 mM KCl, 12 mM EDTA, 0.1 % Triton). Free phosphate was quantified by adding 150 µl of molybdate-malachite green solution (1 mM malachite green, 10 mM ammonium molybdate,1M HCl) and reading the absorbance at 595 nm in a microplate photometer.

### Inositol pyrophosphate synthesis, extraction and quantification

5-InsP7 was synthesized as described (Capolicchio *et al*, 2013; Wang *et al*, 2014). For quantification from cells, InsPPs extraction was performed as described previously (Kim *et al*, 2023), with the following modifications. Briefly, 3 ml of yeast cell culture at OD_600nm_=1 was collected using rapid vacuum-filtration on a polytetrafluoroethylene membrane filter (1.2 μm; Piper Filter GmbH, Germany). After snap freezing in liquid nitrogen, yeast cells on the membrane were resuspended in 400 μL of 1 M perchloric acid and lysed by bead beating (glass beads; 0.25-0.5 mm) for 10 min at 4 °C. After centrifugation at 13,000 rpm for 3 min at 4 °C, the supernatant was transferred into a new tube containing 3 mg of titanium dioxide (TiO_2_) beads (GL Sciences, Japan) which had been pre-washed twice with H_2_O and 1 M perchloric acid. The sample was gently rotated for 15 min at 4 °C. The TiO_2_ beads were collected by centrifugation at 13,000 rpm for 1 min at 4 °C and washed twice using 1 M perchloric acid. After the second washing step, the TiO_2_ beads were resuspended in 300 μL of 3 % (v/v) NH_4_OH and rotated gently at room temperature. After centrifugation at 13,000 rpm for 1 min, the eluants were transferred into a new tube and dried in SpeedVac (Labogene, Denmark) at 42 °C. InsPPs were measured through capillary electrophoresis coupled to mass spectrometry (CE-MS) as described (Qiu *et al*, 2021).

### Quantification of the vacuolar proteome

Vacuoles were prepared as described above. During the purification procedure, a sample of the sedimented spheroplasts was withdrawn before DEAE dextran was added. These withdrawn spheroplasts constitute the “whole cell” extract. After the flotation step, the vacuoles withdrawn from the 4%-0% Ficoll interface were used as the vacuole fraction. For each fraction samples with 100 µg protein were precipitated by adding a final concentration 12.5% TCA for 10 min on ice. The proteins were sedimented (12’000 x g, 5 min, room temperature), the supernatant discarded, and the pellets were washed with ice-cold acetone and centrifuged as above twice. The final pellet was dried and dissolved in reducing SDS sample buffer (10% glycerol, 50 mM Tris-HCl pH 6.8, 2 mM DTT, 2% SDS, 0.002% bromophenol blue).

Samples dissolved in Laemmli buffer (50 mM Tris pH 6.8, 10 mM DTT, 2 % SDS, 0.1 % bromophenol blue, 10 % glycerol) were equalized in concentration by dilution in SP3 buffer (2% SDS, 10 mM DTT, 50 mM Tris, pH 7.5). 150 µg of protein per sample were heated at 95 °C for 5 min and cooled down. Reduced cysteine residues were alkylated by adding iodoacetamide (30 mM final) and incubating for 45 min at room temperature in the dark. Digestion was done by the SP3 method (Hughes *et al*, 2019) using magnetic Sera-Mag Speedbeads (Cytiva 45152105050250, 50 mg/ml). Beads were added at a ratio 10:1 (w:w) to samples, and proteins were precipitated on beads with ethanol (final concentration: 60 %). After 3 washes with 80% ethanol, beads were digested in 50 μl of 100 mM ammonium bicarbonate with 3.0 μg of trypsin (Promega #V5073). After 1 h of incubation at 37 °C, the same amount of trypsin was added to the samples for an additional 1 h of incubation. Supernatants were then recovered, transferred to new tubes, acidified with formic acid (0.5% final concentration), and dried by centrifugal evaporation. To remove traces of SDS, two sample volumes of isopropanol containing 1% TFA were added to the digests, and the samples were desalted on a strong cation exchange (SCX) plate (Oasis MCX; Waters Corp., Milford, MA) by centrifugation. After washing with isopropanol/1%TFA and 2% acetonitrile/0.1% FA, peptides were eluted in 200µl of 80% MeCN, 19% water, 1% (v/v) ammonia, and dried by centrifugal evaporation. Data-dependent LC-MS/MS analyses of samples were carried out on a Fusion Tribrid Orbitrap mass spectrometer (Thermo Fisher Scientific) interfaced through a nano-electrospray ion source to an Ultimate 3000 RSLCnano HPLC system (Dionex). Peptides were separated on a reversed-phase custom packed 45 cm C18 column (75 μm ID, 100Å, Reprosil Pur 1.9 μm particles, Dr. Maisch, Germany) with a 4-90% acetonitrile gradient in 0.1% formic acid at a flow rate of 250 nl/min (total time 140 min). Full MS survey scans were performed at 120’000 resolution. A data-dependent acquisition method controlled by Xcalibur software (Thermo Fisher Scientific) was used that optimized the number of precursors selected (“top speed”) of charge 2+ to 5+ while maintaining a fixed scan cycle of 0.6 s. Peptides were fragmented by higher energy collision dissociation (HCD) with a normalized energy of 32%. The precursor isolation window used was 1.6 Th, and the MS2 scans were done in the ion trap. The m/z of fragmented precursors was then dynamically excluded from selection during 60 s.

Data files were analysed with MaxQuant 1.6.14.0 (Cox & Mann, 2008) incorporating the Andromeda search engine (Cox *et al*, 2011). Cysteine carbamidomethylation was selected as fixed modification while methionine oxidation and protein N-terminal acetylation were specified as variable modifications. The sequence databases used for searching were the S. cerevisiae reference proteome based on the UniProt database (https://www.uniprot.org, version of June 6th, 2021, containing 6050 sequences), and a “contaminant” database containing the most usual environmental contaminants and enzymes used for digestion (keratins, trypsin, etc). Mass tolerance was 4.5 ppm on precursors (after recalibration) and 20 ppm on MS/MS fragments. Both peptide and protein identifications were filtered at 1% FDR relative to hits against a decoy database built by reversing protein sequences. The match between runs feature was enabled.

## Data availability

Source data from the proteomic analysis have been deposited at the PRIDE database under the identifier PXD060102.

## Acknowledgements

Mass spectrometry-based proteomics work was performed by the Protein Analysis Facility of the Faculty of Biology and Medicine, University of Lausanne. This study was supported by grants from the SNSF (320030-228119, 31003A_179306 and 310030_204713) and ERC (788442) to AM, by the HFSP (LT000588/2019) to GDK, by the Deutsche Forschungsgemeinschaft (CIBSS, EXC-2189, Project ID 390939984, to HJJ) and the Volkswagen Foundation (VW Momentum Grant 98604 to HJJ).

## Author contributions

AM conceived the study, analysed data and wrote the paper. SB performed in vitro and in vivo experiments with yeast. GDK, LM, GL, and HJJ analysed inositol pyrophosphates. All authors assembled and corrected the manuscript.

## Competing interests

None

## Notes

https://www.ebi.ac.uk/pride/archive/projects/PXD060102

### Competing Interest Statement

The authors have declared no competing interest.

### Summary of Updates

Titration of Pi sensitivity of polyPases Sample picture of isolated vacuoles

